# A Novel Single-Fish Assay for Ethanol Self-Administration in Zebrafish

**DOI:** 10.64898/2026.05.26.727885

**Authors:** Léa Morneau, Léa Gagné, Randall T. Peterson, Gabriel D. Bossé

## Abstract

Alcohol use disorder (AUD) is a significant public health concern. In Canada, about 18% of individuals aged 15 or older will meet the clinical criteria for AUD at some point in their lives (CAMH, 2023). Treatment options for AUD are limited, and the high relapse rates highlight the urgent need for innovative methods to study and address AUD.

Zebrafish (Danio rerio) is an emerging model for exploring the neurobiological impacts of alcohol. Previous studies have demonstrated that zebrafish respond to the rewarding effects of alcohol, but most research methods rely on passive administration, such as immersion, which does not reflect the typical routes of alcohol intake in humans.

We previously showed that zebrafish can learn to self-administer drugs of abuse in small groups and conditioned animals are displaying key features of substance abuse disorders. However, group-based conditioning limits our understanding of individual drug preference and intake profile.

In this study, we improved upon our previous design by establishing an individual self-administration protocol to measure voluntary alcohol intake and model alcohol use disorder. In this novel assay, individual adult fish learn to discriminate between two zones to self-administer a 5% ethanol solution. Moreover, animals conditioned in this assay can perform progressive ratio and display signs of withdrawal upon cessation of ethanol intake. These results suggest zebrafish can develop ethanol abuse-like behaviour, providing a powerful platform to study genetic predisposition and screen for therapeutic compounds.

## 1. Introduction

A recent study revealed that alcohol is the most abused substance in Canada^1^. It is estimated that over 90% of the population have consumed alcohol at least once, and 18% of Canadians over the age of 15 have met the criteria for alcohol use disorder (AUD) at some point in their lives (CAMH, 2023). The World Health Organization also reported that alcohol is responsible for 2.6 million deaths per year worldwide, representing 4.7% of all deaths. In addition to the human cost, the United States, excessive alcohol use is associated with over $240 billions in economic cost^2,3^.

AUD is a medical condition that can be described as an impaired ability to stop using, regardless of adverse consequences^4^.

Alcohol use disorder arises from the interaction of several neurobiological processes, including acute reinforcement, neuroadaptation, withdrawal, craving, and relapse^5–7^. Ethanol acts on multiple neurotransmitter systems, enhancing GABAergic inhibition, modulating glutamatergic NMDA receptor signaling, and engaging mesolimbic dopamine pathways that contribute to reward and reinforcement^5,6,8^. With repeated exposure, compensatory adaptations in these systems can produce tolerance and withdrawal, including anxiety, hyperexcitability, sleep disruption, and negative affective states^9^. These withdrawal symptoms are clinically important because they can drive continued alcohol intake and relapse, even when individuals are motivated to reduce consumption^10^. Current pharmacological treatments for AUD partly target these mechanisms: for example, naltrexone reduces opioid-mediated reward signaling, acamprosate is thought to stabilize glutamatergic/GABAergic imbalance during abstinence, and disulfiram produces aversive physiological effects after alcohol consumption^11,12^. However, treatment responses remain heterogeneous, suggesting that our understanding of the individual behavioural and neurobiological signatures that predict intake, withdrawal severity, and relapse vulnerability remains incomplete. Moreover, even after successful cessation, the relapse rate remains high, with 50% of patients resuming consumption within the first 90 days following treatment^13^. Consequently, novel approaches are needed to improve our understanding of the neurobiological mechanisms driving alcohol use disorder to uncover novel therapeutic targets.

Over the last twenty years, zebrafish have become an innovative model for neurobiological research and, more recently, for exploring the neurobiology of substance use disorders^14–17^. Previous research has shown that zebrafish display robust behavioural and physiological responses to a broad spectrum of drugs of abuse, including psychostimulants, alcohol, nicotine, and opioids, in paradigms commonly used to assess drug-induced reward, tolerance, and sensitization in other model^18–25^. Moreover, key neuronal circuits like dopaminergic, glutamatergic, and opioid systems are highly conserved, and the pharmacodynamic properties of drugs of abuse are similar, which makes them valuable for translational research on substance abuse^15,26,27^. Taken together, these findings support the utility of zebrafish as a tractable and cost-effective model for investigating the mechanisms underlying substance abuse.

Preclinical alcohol research relies on several complementary paradigms, each capturing different aspects of alcohol exposure and AUD-related biology. In zebrafish and other aquatic models, repeated ethanol immersion is commonly used to model acute intoxication, repeated binge-like exposure, tolerance, withdrawal-like phenotypes, and circuit-level consequences of alcohol exposure^28–30^. These approaches are highly valuable because they allow precise control over exposure concentration and timing, are scalable, and can be combined with behavioural, imaging, or molecular endpoints. However, immersion-based paradigms primarily model passive alcohol exposure rather than voluntary alcohol seeking or individual differences in consumption. As a result, they are well-suited to studying the consequences of alcohol intoxication and withdrawal but are less able to capture why some individuals escalate intake, persist in alcohol seeking, or show greater relapse vulnerability. Rodent paradigms such as two-bottle choice, drinking-in-the-dark, operant self-administration, and vapour exposure address some of these limitations by modelling voluntary consumption, binge-like intake, reinforcement, dependence, or relapse-like behaviour^31–34^. Nevertheless, these models can have lower throughput, be more costly, and less suited to systematic screening across developmental stages, genetic backgrounds, or treatment conditions. Therefore, there is a need for scalable models that preserve key strengths of active ethanol exposure while adding quantitative measures of voluntary intake, motivation, and individual behavioural trajectories. Such an approach would allow alcohol-related phenotypes to be studied not only at the group level, but also as individual signatures of vulnerability and resilience.

The gold standard paradigm to model substance abuse disorders involves operant conditioning, such as self-administration, where animals must actively perform a task to receive a drug dose. These assays provide a closer representation of human consumption and allow modelling of various aspects of the substance abuse process, from initial consumption to escalation and more complex consumption patterns, and facilitate the study of neurobiological effects^35–37^. Critically, the neurobiological impacts induced by repeated drug use differ between passive and active consumption^36^; thus, the development of robust self-administration paradigms is essential to model AUD. There is evidence that zebrafish can actively seek and consume drugs of abuse; for instance, mixing ethanol with gelatin will lead to increased consumption compared to just gelatin^39^ and adult fish will voluntarily swim in an active zone to receive ethanol during a single 28-minute session^40,41^. Although these are evidence of voluntary ethanol consumption, these methods are less effective at assessing whether animals will actively seek out ethanol, increase their intake over time, or exhibit individual behavioural patterns linked to AUD development.

Considering these limitations, our group has developed a zebrafish self-administration assay to assess substance intake and drug-directed behaviour at the individual level. This assay provides a scalable framework for quantifying voluntary drug access, intake trajectories, and behavioural phenotypes that may be difficult to resolve with passive exposure or feeding-based paradigms alone. This assay is based on a paradigm previously developed in our group to study opioid self-administration assay, where a small group of adult zebrafish were conditioned to swim to an active platform to receive a dose of opioids^25^. Fish conditioned in the arena displayed several hallmarks of substance abuse, such as seeking despite adverse consequences, increased motivation for the drug, and they exhibited signs of stress following drug cessation. However, in this set-up, it was not possible to follow the consumption patterns of individual fish and given that zebrafish are social animals that move in shoals^42^, group dynamics could also influence consumption.

To overcome these limitations, we modified our assay and developed an individual ethanol self-administration assay in adult zebrafish. In this test, fish can trigger the administration of an ethanol solution by actively swimming in a specific zone arena. Fish trained to self-administer ethanol showed an increased preference for the drug-paired zone across training days. These animals also performed progressive ratio conditioning and displayed signs of withdrawal. As such, this new paradigm is the first application of operant conditioning to individual zebrafish, allowing direct measurement of consumption over time and enabling identification of individuals with greater vulnerability and increased ethanol preference. The development of this novel test will lead to better modelling of AUD in zebrafish to further develop an understanding of the biological mechanisms underlying drug seeking and open the door for the development of more complex behaviours related to substance abuse disorder, such as incubation of craving and relapse studies.

## 2. Methods

### 2.1 Animal housing

Adult zebrafish (*Danio rerio*) of the wild-type Tüpfel long fin (WT-TL) line were maintained in the fish facility at 28−29°C with a 14/10 h light/dark cycle. Fish were raised following these procedures. Embryos were obtained from multiple mating pairs, pooled and randomly assigned to treatment groups. Animals were raised in E3 solution (NaCl 4.97 mM, KCl 0.179 mM, CaCl2 0.33 mM, MgCl2 0.40 mM, Hepes 20 mM) at a pH of 7.2 in 100 mm x 25 mm petri dishes (Avantor Science Central, USA) at a density of 1 embryo/ mL and kept at 28.5°C with a 14/10 hours light/dark cycle. At 3 dpf, chorion debris were removed.

From 5 to 10 dpf larvae were fed daily with 30 μL/larvae of eukaryotic *Tetrahymena termophila. T. thermophila* cultured in an SPP medium enriched with peptone protease, and Fe3+ (2% proteose peptone, 0.1% yeast extract, 0.2% glucose, 33 μM FeCl3, 250 μg/mL Streptomycin/penicillin, 0.25μg/mL Amphotericin B) until their concentration reached a plateau. An 11 mL volume of SPP medium was inoculated with 1 mL of *T. thermophila* culture at maximum confluence. Before use, *T. thermophila* were washed in E3 medium from their SPP medium to remove antibiotics, using three centrifugations (2 min, 0.8 g).

At 10 dpf, larvae were transferred into 1.1 L recirculating tanks filled with deionized water supplemented with Instant Ocean Sea Salt (Tropic Marin, Wartenberg, Germany) and NaHCO₃ to maintain a conductivity of 700–1000 μS and a pH of 7.2–7.4. Larvae were housed in groups of 25–30 individuals per tank and maintained at 28.5°C under a 14:10 light-dark cycle. Water temperature, conductivity, and ammonia levels were monitored daily. From 10 dpf onward, larvae were fed daily with 24-hour-hatched *Artemia nauplii* until the end of the experiment. The amount of food was based on intake for the first 10 minutes to ensure it was not limited in quantity, as described in Caperaa *et al.* 2025^43^.

All experiments described in this study were approved and performed in compliance with the *Comité de Protection des Animaux* (CPAUL) of Université Laval. All experiments were performed on a mixed population of male and female fish (50-50% ratio).

### 2.2 Experimental apparatus

#### 2.2.1 Design of the testing arena

The conditioning arena consists of a 3D-printed plastic Z shaped arena (Figure 1A) connected to a pump (Supreme Aqua-Mag, Thatfishplace, USA) to generate a continuous recirculating flow of water (Figure 1A) to prevent drug build up. In this design, the extremities of the arenas consist of the drug-paired and inactive zones. The arena is illuminated with a warm white light source (2700 K, 40 W, CFL bulb, McMasterCarr, USA), providing enough light to allow the fish to identify the different zones without affecting behaviour. Infrared cameras (PiNoir camera, Adafruit, USA) are installed over the arena and are connected to a mini-computer Raspberry Pi3 (Adafruit, USA) to monitor movement in each zone. The computer connected to the camera also controls a peristaltic 12 V pump (Adafruit, USA). A small silicone tube is attached to the side of the active zone to enable direct drug delivery.

**Figure 1.**
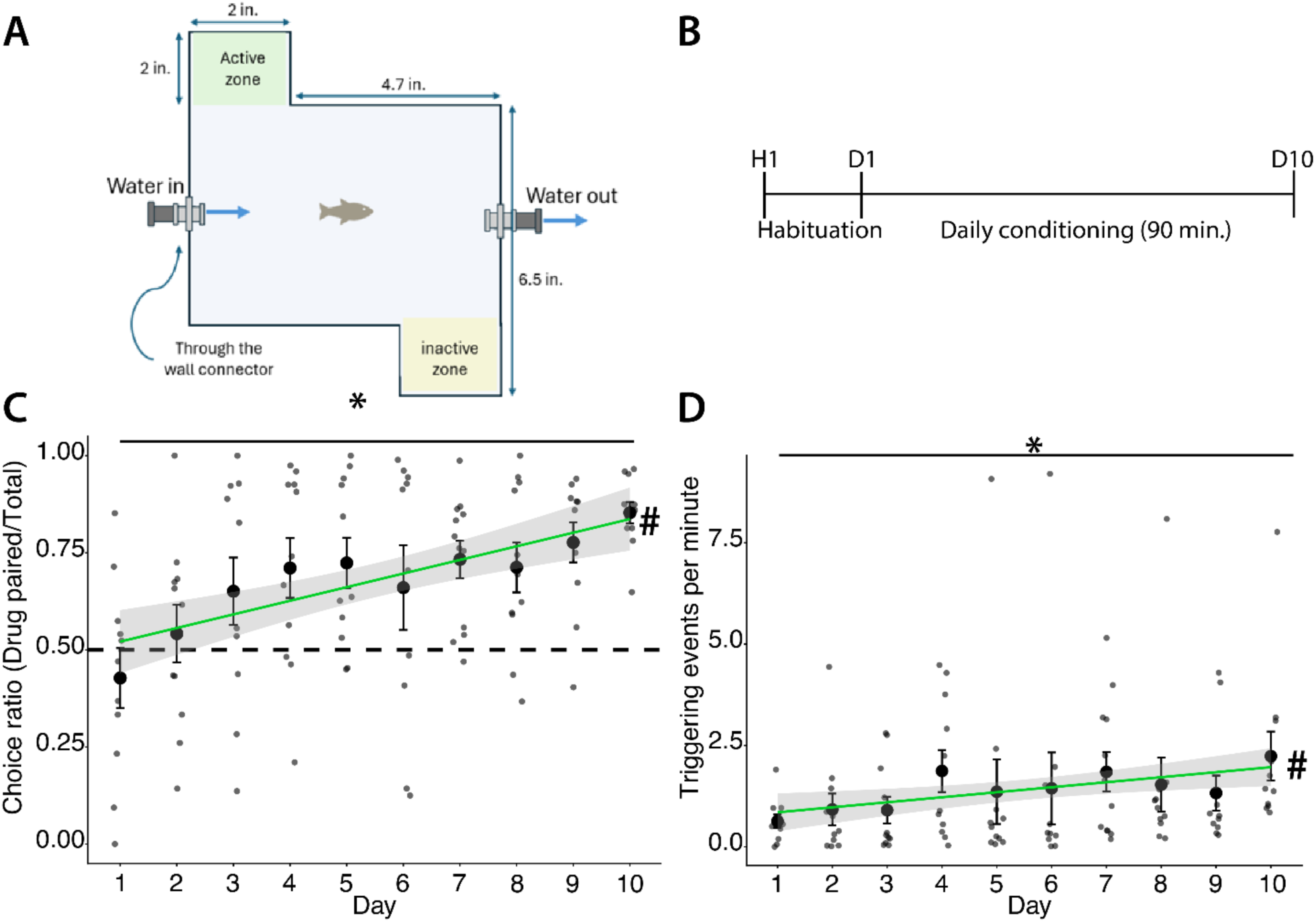
Individual zebrafish ethanol self-administration conditioning. The training starts with 2 days of habituation (H1), followed by 10 days of daily conditioning. **A:** Layout of the conditioning arena. **B:** Overview of the conditioning timeline. **C:** A choice ratio was calculated for each conditioning session; this ratio represents the number of drug-paired triggering events divided by the total number of events. A ratio above 0.5 indicated a large number of drug-paired visits. The average Choice ratio increases across the days of conditioning. \**p*-value < 0.05 calculated using Student’s t-test comparing the average on Day 10 vs Day 1. # *p*-value < 0.05. The day-dependent trend was assessed using simple linear regression (Choice ∼ Day), which showed a significant positive association between testing day and the choice ratio (β = 0.035 ± 0.008 SE, p = 1.17e-05). **D:** The rate of activation (Triggering events per minute) increases over the training session. \**p*-value < 0.05 calculated using Student’s t-test comparing the average on Day 10 vs Day 1. # *p*-value < 0.05. The day-dependent trend was assessed using simple linear regression (Rate ∼ Day), which showed a significant positive association between testing day and the number of triggering events per minute (β = 0.12 ± 0.06 SE, p = 0.0409). Larger dots represent the group average, while the small ones represent an individual. Error bars indicate the standard error of the mean (S.E.M.). n=11. Experiment performed in a within-subject design.

#### 2.2.2. Development of the coding script

Our assay runs on a Raspberry Pi3 with the latest Raspbian OS, controlled by a custom Python 3 script. The script detects fish movement within a zone by comparing the current frame’s pixels to the average of the previous frames. A movement event is triggered if the pixel difference exceeds a manually set threshold, which was adjusted for each experiment, to prevent the circulating water from triggering the pump. The threshold was set manually before each experiment to account for small variations in water flow or zone size as they are manually selected. The code records the elapsed time, saves an image of the frame with detected motion, and logs the number of triggers in each zone. Additionally, a short delay after pump activation ensures that the animals exit the zone before the pump can trigger again. Each trigger delivered 250 mL of ethanol 5 % directly in the drug-paired zone.

### 2.3. Animal conditioning

#### 2.3.1. Habituation

To minimize stress during training, zebrafish are initially exposed to the arena in groups of three. On the first day, they are placed together in the arena for 90 minutes. Animals are free to explore the arena, and no reward is offered. The next day, they are again placed in groups of three for 90 minutes and now ethanol was released upon entering the active zone so fish can learn that a reward is offered in the arena.

#### 2.3.2. Ethanol self-administration conditioning

After habituation sessions, each animal was trained for 90 minutes daily. Since zebrafish are social animals, to avoid stress from social isolation, animals were kept in small groups of 3 to 6 between sessions. To prevent time-of-day bias, animals were conditioned in a random order each day. During each training session, individual fish could swim freely in the arena, and when entering the drug-paired zone, they received a 5% ethanol dose diluted in water from the main system. The conditioning used a fixed-ratio schedule, where one triggering event resulted in one dose of the drug. Continuous water flow prevented ethanol saturation and prompted the fish to reactivate the pump by re-entering the active zone for a new dose.

#### 2.3.3 Progressive ratio conditioning

The Python script was modified to implement a progressive ratio paradigm by increasing the number of visits to the drug-paired zone required to trigger the delivery of one dose of ethanol by 1 following each triggering event within the same session. As a result, the fish must enter the drug-paired zone multiple times to receive a single dose, with each subsequent delivery requiring more visits. In our setup, the breakpoint was defined as the highest number of triggering events per dose delivered achieved during each daily session.

### 2.4. Dopamine receptor inhibitor treatment

Conditioned animals were exposed to 10 µM SCH-23390 hydrochloride for 60 minutes, as previously performed^25^. The drug was directly transferred to the fish water.

### 2.5. Stress-induced withdrawal measurement

The novel tank assay was conducted using the *Zantik LT* system and a 129 x 360 mm tank, also from Zantiks, UK, filled with water from the main system. The test took place between 9am and 11am, with each fish being recorded for 12 minutes. Animal movement was tracked using *Zantik* software, which defined three zones; top, middle, and bottom, to measure exploration. Before starting, each fish was placed in the center of the tank. The system automatically tracked movement, recording the animal’s position (x,y,z). The software then calculated freezing time and distance travelled within each zone based on these coordinates.

Raw data generated by the *Zantiks system* were analyzed using custom R scripts. Time was analyzed on log10-transformed values using a linear model with condition as a fixed effect. Planned comparisons of Day2 and Day6 versus Control were performed using estimated marginal means with Dunnett adjustment. Data were log10-transformed to reduce right skewness, improve normality, and stabilize variance.

### 2.6. Chemical treatment

The following chemicals were used in this paper: ethanol at 5% (volume/volume) (Les alcools de commerce, Canada) and SCH-23390 hydrochloride (Tocris, UK).

### 2.7 Data visualization

All box plots were generated using R graphics programming and the *ggplot* module.

### 2.8 Statistical analysis

A *P* value of less than 0.05 was considered significant. The specific statistical analysis for each Figure panel is listed in its respective figure legend.

## 3. Results

### 3.1 Ethanol self-administration conditioning

We previously demonstrated that a small group of adult zebrafish can be conditioned to self-administer various opioids^25,44^. Since the training involved multiple animals, individual consumption could not be monitored, and social interaction might have influenced the training effectiveness. One limitation of the group-based protocol was the arena’s size, which caused stress when individual fish were introduced. We thus designed a smaller arena based on an earlier model that used a single-fish training task (Figure 1A). While our previous studies conditioned animals with opioids, we opted to develop this new paradigm with ethanol, as research indicates it has a strong rewarding effect and is highly attractive to adult zebrafish^39^.

This new paradigm is based on the same core principle as our earlier group self-administration model. The key difference is that we created a custom 3D-printed Z-shaped arena, where each end is designated as either the drug-paired zone or the inactive zone. A continuous water flow ensures that drugs do not accumulate. In this paradigm, individual adult fish were trained to swim towards the drug-associated zone to receive an ethanol dose. During each training session, we counted the number of triggering events in both the drug-paired and inactive zones. We then calculated a choice ratio by dividing the number of drug-paired triggers by the total triggers. A ratio above 0.5 indicated that the fish preferred the drug-paired zone.

Individual fish underwent conditioning over ten days (Figure 1B). By day 1, some showed a clear preference for the drug-paired area, though the average choice ratio was about 0.5. Over time, the choice ratio increased significantly, reaching 0.8 by day 10 (Figure 1C). We also monitored the number of visits to the drug-paired zone per minute as a proxy for consumption. Throughout the protocol, these visits increased significantly, by 237% from day 1 to day 10 (Figure 1D). These results suggest that individual fish can learn to self-administer ethanol within 10 days and progressively increase their consumption.

### 3.2 Conditioned animals show learning flexibility

To ensure the animals understood the task and were motivated by ethanol, we conditioned a new group for 5 days using the same approach previously described. Then, over the next 5 days, we moved the drug-delivery tube to the inactive zone, thereby reversing the zones (Figure 2A). On Day 6, the first day of the switch, animals still favoured the original drug-associated zone, as shown by their choice ratio below 0.5. However, from Day 7 onward, they adapted their behaviour and learned to visit the new drug-paired zone (Figure 2B). This demonstrates that individual animals are motivated by alcohol, understand the task and showed behavioural flexibility when faced with changing parameters.

**Figure 2.**
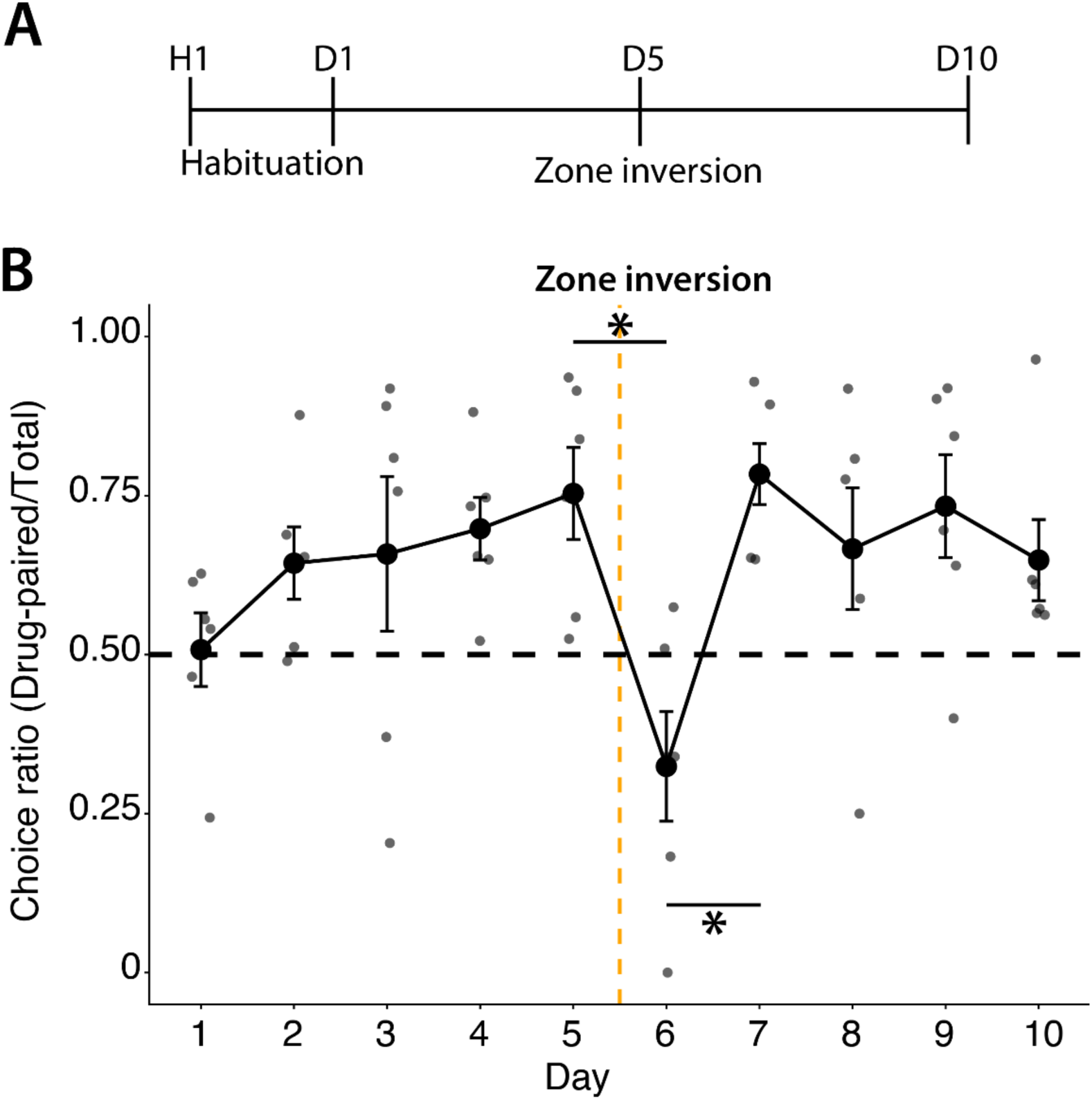
Animals adapt to zone inversion. **A:** Overview of the reverse-conditioning timeline. **B:** After the first 5 days of conditioning, the location of the drug-paired zone was changed. Animals were able to identify the correct drug-paired zone by the second day of reversal learning. Session number was indexed within phase, and behaviour was analyzed using a linear mixed-effects model with fish identity as a random effect. There is a significant reduction in the choice ratio between the first day of reversal (Day 6) and both the last day of the initial phase (Day 5) and the second day of reversal (Day 7), but phase-matched comparisons showed that performance did not differ significantly between the initial and reversed contingencies from session 2 onward, suggesting rapid adaptation after the switch. Larger dots represent the group average, while the small ones represent an individual. \**p*-value < 0.05 calculated using linear mixed-effects model followed by Holm-adjusted post hoc pairwise comparisons. Error bars indicate the standard error of the mean (S.E.M.). n=6. Experiment performed in a within-subject design.

### 3.3 Successful progressive ratio conditioning

Our initial training employed a fixed-ratio schedule in which one triggering event resulted in one dose of drug. In other animal models, the progressive ratio has been suggested as a more reliable indicator of motivation than fixed-ratio training^45–47^. During progressive ratio training, animals must perform an increasing number of actions to obtain each dose. This contrasts with fixed-ratio conditioning, in which each action directly leads to drug delivery. For instance, an animal might need to press a lever to receive a dose of drug, with the required presses escalating until the animal reaches its limit, known as the breakpoint. Usually, the progressive ratio is tested on animals that have had prior fixed ratio training. This approach also helps us confirm that the animals did not simply develop a habitual swimming pattern or a basic zone preference, as the fish should lose interest if no ethanol is delivered or show no adaptation to the progressive ratio conditioning.

We therefore chose to perform progressive ratio training to evaluate the motivation of ethanol-conditioned animals to consume ethanol (Figure 3A). We conducted progressive ratio conditioning over five consecutive days and observed that conditioned fish increased their total visits each day, reaching a higher breakpoint (Figure 2B-C). One individual even reached a breakpoint of 100 visits to receive a single dose. This rising breakpoint also suggests that the amount of ethanol delivered continued to increase as progressive ratio training progressed.

**Figure 3.**
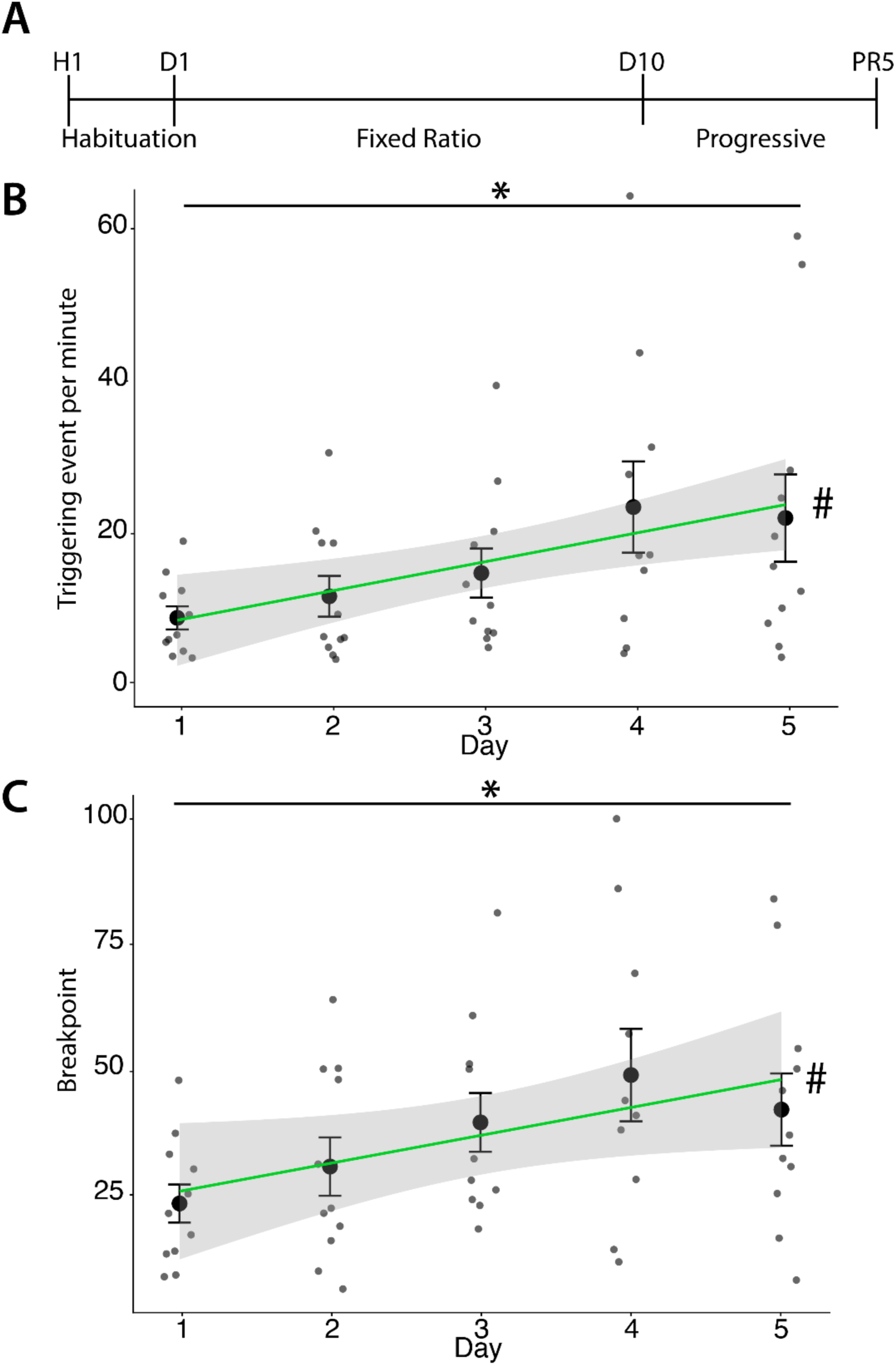
Conditioned animals can perform progressive ratio conditioning. **A:** Overview of the conditioning timeline. **B:** During progressive ratio conditioning, the number of triggering events required to receive a dose of ethanol was increased by 1 following each drug delivery. We observed a significant increase in the number of visits per minute across the progressive ratio training sessions. \**p*-value < 0.05 calculated using Student’s t-test comparing the average on Day 5 vs Day 1. # *p*-value < 0.05 The day-dependent trend was assessed by simple linear regression (Triggering events per minute ∼ Day), showing a significant positive association between testing day and breakpoint (β = 3.811 ± 1.292 SE, p = 0.00477). **C:** The breakpoint was defined as the highest number of triggering events per drug delivery completed by an individual animal. The breakpoint was significantly higher on Day 5 than on the first day of progressive ratio conditioning, and the average breakpoint increased over the session. \**p*-value < 0.05 calculated using Student’s t-test comparing the average on Day 5 vs Day 1. # *p*-value < 0.05 represents the day-dependent trend was assessed by simple linear regression (Breakpoint∼ Day), showing a significant positive association between testing day and breakpoint (β = 5.51 ± 2.03 SE, p = 0.009). Larger dots represent the group average, while the small ones represent an individual. Error bars indicate the standard error of the mean (S.E.M.). n=11. Experiment performed in a within-subject design.

The increase in the total number of triggering events detected with the progressive ratio indicates that ethanol-conditioned fish are willing to repeatedly perform the required action to obtain a dose. This result suggests that pump activation is not simply a conditioned place preference, but rather an active self-administration of the drug.

### 3.4 Ethanol self-administration is driven by dopaminergic signalling

Ethanol affects the brain partly by enhancing GABAergic signalling at certain receptors and by altering inhibitory control over dopamine neurons, thereby increasing dopamine release in reward circuits^6^. Since dopamine is important for developing substance abuse disorder and serves as a key neurotransmitter in the reward system, we investigated the role of dopaminergic signalling in ethanol self-administration. We used the dopamine D1 receptor antagonist SCH-23390, which prior studies have shown to decrease motivated behaviours, including drug self-administration, in zebrafish and other models^19,48^ without affecting general locomotion^25^. Initially, individual fish were conditioned over 4 days using a fixed-ratio protocol (Figure 4A). On the fifth day, animals were treated 10 µM SCH-23390 for 1 hour before their training session. Notably, this treatment with the D1 receptor antagonist significantly lowered the choice ratio (Figure 4B), supporting that ethanol self-administration in adult zebrafish relies on dopaminergic signalling.

**Figure 4.**
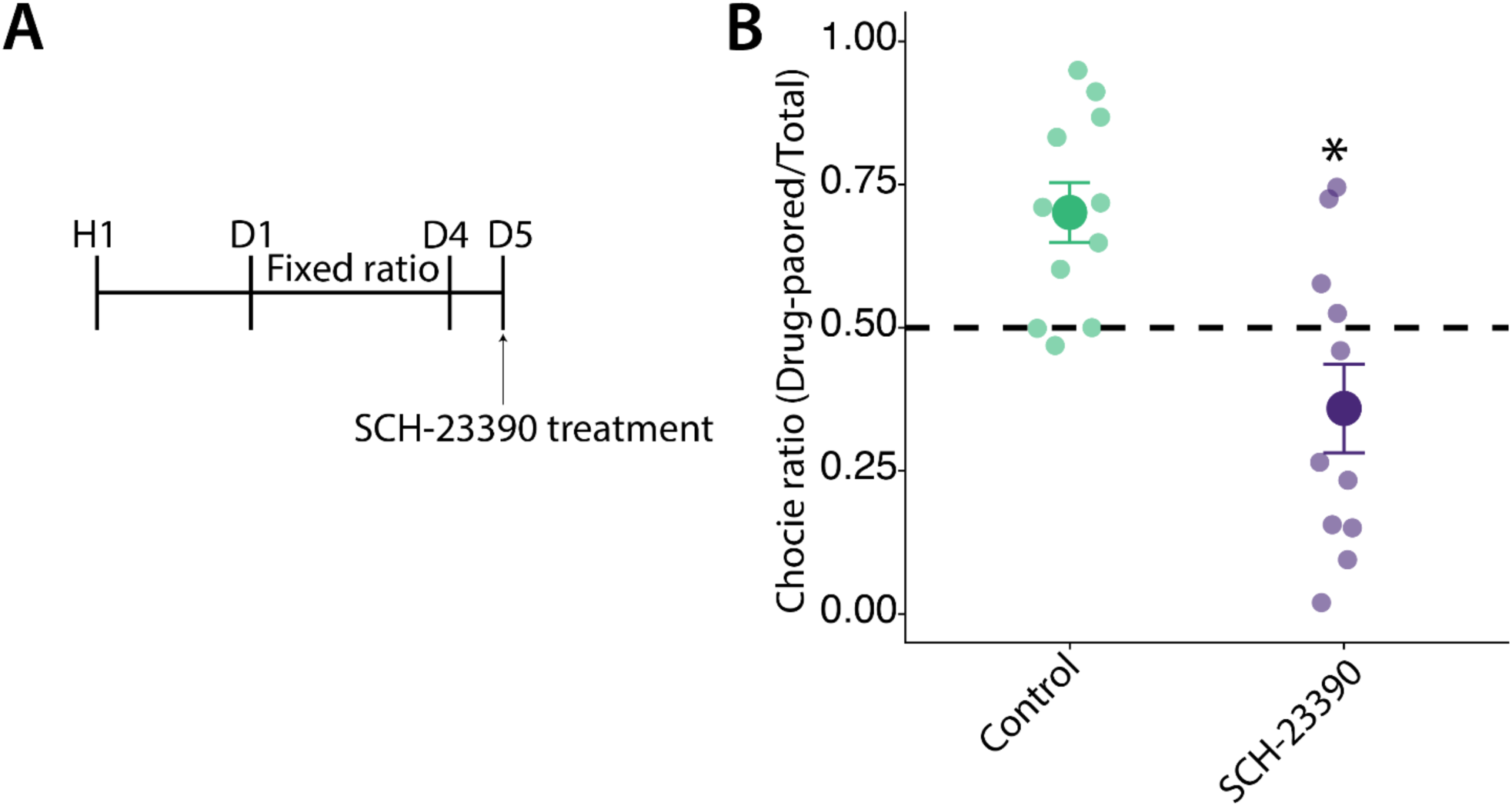
The choice ratio is dopamine-driven. **A:** Timeline of the experiment. Animals were first conditioned for 4 days in the arena. Before the 5^th^ session, they were exposed to SCH-23390 (10 µM for 60 minutes). **B:** SCH-23390 treatment reduces the choice ratio. \**p*-value < 0.05 calculated using Student’s t-test comparing the average. The large dot represents the group average, while the smaller dots represent individual choice ratios. Error bars represent the S.EM. n=11. Experiment performed in a within-subject design.

### 3.5 Development of withdrawal symptoms following drug discontinuation

The development of withdrawal symptoms after stopping drug use is a key feature of substance abuse disorder. These symptoms include both intense physical and psychological effects, such as increased anxiety and stress levels in both humans and animal models. These symptoms reflect neurobiological adaptations to repeated drug consumption and underlie the continued drug seeking seen in patients with substance use disorders^49^. In zebrafish, previous studies have demonstrated that withdrawal to different drugs can cause profound anxiety/stress^50^ that can manifest as reduced exploration behaviour, increased erratic movements, changes in freezing behaviour, and changes in locomotion. In a previous study, Mathur and Guo demonstrated that ethanol withdrawal in zebrafish peaked a few days after drug discontinuation^51^. Therefore, we examined the behaviour of conditioned fish 2 and 6 days after their last self-administered ethanol dose, using the well-established novel tank assay to evaluate stress and anxiety in zebrafish (Figure 5A).

**Figure 5.**
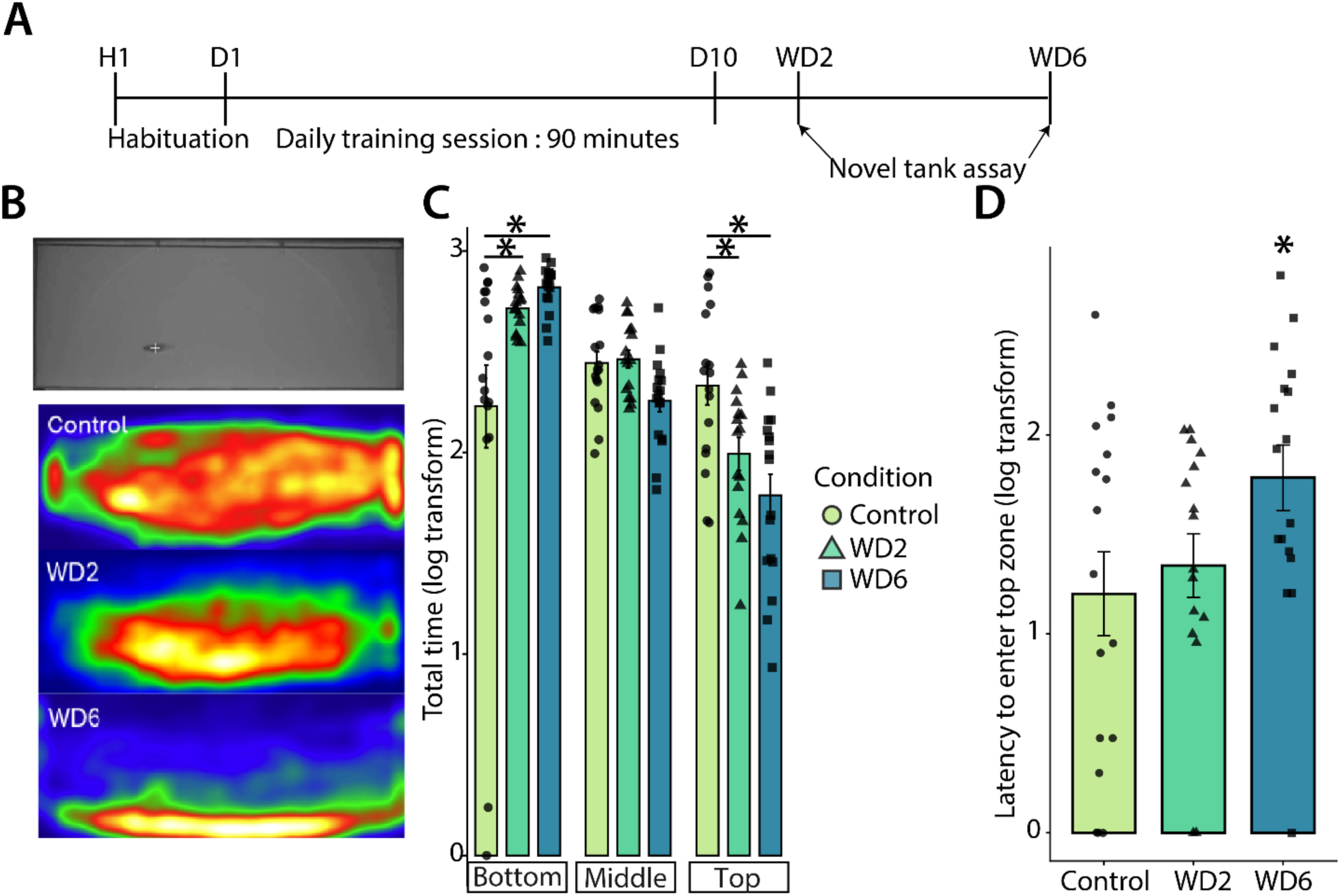
Drug discontinuation triggers an increase in stress. **A:** Timeline of the experiment. D=Day of ethanol conditioning, WD = withdrawal day. **B:** Representative heat map of the location viewed for the side of one fish in the novel tank assay for each condition. **C:** Average time spent in each of the arena’s zones. Animals at WD2 and WD6 spent less time in the top zone and more time at the bottom compared to control animals. WD6 to control. **D:** The latency to enter the top zone was greater in the WD6 animals compared to controls. \**p*-value < 0.05. Statistical analysis was performed on log10-transformed time values using a linear model followed by Dunnett-adjusted comparisons versus Control. n=16. Experiment performed in a between-subject design.

In this assay, individual fish are introduced into a new tank to measure their exploration behaviour by analyzing the total time spent swimming at the bottom of the tank versus in the top and middle areas. Stressed animals have been shown to exhibit reduced exploration of a novel environment and avoid the surface of the water^52^. As expected, zebrafish in withdrawal showed less exploratory behaviour, spending more time in the bottom zone than the top at both 2 and 6 days following drug discontinuation (Figure 5B-C). We also measured the latency to reach the top zone which was significantly longer at day 6. (Figure 5D) The results suggest that conditioned fish in withdrawal display signs of stress and anxiety as demonstrated by reduced exploration activity. This increase in stress during withdrawal further supports the idea that conditioned zebrafish develop addiction-like behaviour to ethanol.

## 4. Discussion

This study presents the development and validation of a new self-administration assay for adult zebrafish, offering a novel method to examine voluntary alcohol consumption and reward-driven learning. The conditioned animals showed behaviours similar to operant responding and features of substance abuse disorder. Overall, these results support the use of adult zebrafish as a model for ethanol self-administration research.

### 4.1 Developing substance abuse-relevant assays in zebrafish

Alcohol use remains a major societal challenge with important health and economic impacts; treatment options remain limited, and novel approaches are needed to study neurobiological pathways driving the development and sustain drug use. Zebrafish has emerged as an innovative model to study substance abuse neurobiology. However, many existing zebrafish drug exposure paradigms rely on non-contingent administration or conditioned preference, which do not directly assess voluntary drug intake. Previous work by *Sterling et al (2014)*^39^ and *Raine et al (2025)*^41^ demonstrated that adult zebrafish actively consume a mixture of agar and ethanol and choose to swim to a zone to consume ethanol; these studies give an insight into alcohol preference, however it doesn’t address classic addiction-like behaviours

We previously developed a group-based opioid self-administration conditioning paradigm in which zebrafish were trained to swim over an active platform to receive an opioid dose. Using this approach, conditioned animals exhibited several addiction-relevant behaviours. However, the group-based design did not permit precise tracking of individual drug intake, and social dynamics could influence drug-seeking behaviour, introducing potential confounding factors. To address these limitations, we developed an individual-based drug self-administration assay that allows direct measurement of drug-seeking behaviour at the single-animal level.

The development of robust substance abuse–relevant behavioral paradigms, such as drug self-administration, will enable researchers to more fully leverage the strengths of the zebrafish model to advance our understanding of the neurobiology of substance use disorders (SUDs). Notably, genetic factors are estimated to account for up to 60% of individual vulnerability to developing SUDs^53,54^, yet the underlying mechanisms remain incompletely understood. Although genome-wide association studies have identified numerous candidate genes, the causal roles of many of these variants remain largely untested in vivo.

Zebrafish offer a powerful platform to address this gap, as straightforward and efficient gene-editing approaches, including CRISPR–Cas9, permit the rapid generation of targeted mutant lines. This capability allows systematic investigation of gene function in drug-seeking behavior and addiction-relevant phenotypes. Indeed, prior studies in zebrafish have shown that manipulating specific genes (e.g., *ccser1*) alters ethanol preference, supporting the translational relevance of this model^40^. In addition to genetic approaches, the scalability of zebrafish and compatibility with precise pharmacological manipulation facilitate unbiased small-molecule screening to identify compounds that reduce drug consumption^44^. Such screens have the potential to reveal novel biological pathways involved in substance abuse and to accelerate the discovery of new therapeutic strategies.

### 4.2 Individual conditioning

Building on prior studies showing that adult zebrafish voluntarily consume ethanol, we hypothesized that ethanol functions as a robust reinforcer and is particularly well-suited for the development and optimization of an individual-based conditioning paradigm. We thus adapted a new arena for individual fish conditioning. Critically, animals could only receive ethanol by actively swimming to the drug-paired zone, as the continuous water flow prevented drug accumulation in the tank. The custom script also incorporated a brief time-out after each trigger, requiring animals to leave the drug-paired zone before another dose could be given. One of the main strengths of this assay is its capacity to differentiate reinforced responses from general locomotor or exploratory behaviours. The rising choice ratio indicates that animals quickly learn to associate the correct zone with drug delivery, consistently swimming towards the drug-paired area. This behaviour suggests that their actions are guided by learned associations rather than by increased exploration or general activity. Moreover, because animals are free to explore the entire arena but must actively visit the drug-paired zone to obtain a dose, drug delivery is contingent on operant-like spatial action rather than being passively imposed; we consider these features consistent with self-administration paradigms established in other animal models.

### 4.3 Evidence of self-administration and hallmarks of substance abuse disorder

One of the defining hallmarks of drug addiction is the escalation of drug intake over time, reflecting increased motivation and reinforcement rather than passive exposure. In our paradigm, we observed a progressive increase in the number of triggering events across conditioning sessions, consistent with escalation phenomena reported in established self-administration models, suggesting that animals learned the action–outcome contingency required to obtain the drug. To further strengthen the paradigm’s self-administration nature, we assessed behavioural flexibility by reversing the location of the active zone. Notably, fish conditioned for five days rapidly adjusted their behaviour following zone swapping, preferentially targeting the new drug-paired zone. This finding further supports the conclusion that drug-seeking behaviour was guided by learned, goal-directed decision-making rather than a rigid or habitual response; the emergence of this pattern therefore supports both task acquisition and the reinforcing properties of ethanol within the assay. This also further demonstrates that adult zebrafish exhibit adaptive behavioural flexibility within this self-administration framework.

To further investigate motivation-driven behaviour, we adapted a progressive ratio conditioning paradigm, which is widely regarded as a sensitive and robust readout of motivational strength in self-administration studies. In progressive ratio schedules, the response requirement to obtain a drug dose increases systematically, thereby providing a quantitative measure of the effort a subject is willing to exert for reinforcement. The breakpoint, defined as the final ratio completed before responding ceases, is a key parameter commonly used to estimate and compare maximal motivational drive across subjects.

In the present assay, performance under the progressive ratio schedule indicates that zebrafish progressively increased the number of triggering responses required to compensate for reduced drug availability, thereby raising their breakpoint and demonstrating a clear link between operant action and drug delivery. The ability of conditioned fish to sustain responding under an intra-session progressive ratio schedule strongly supports the interpretation that triggering events reflect voluntary, motivation-driven self-administration. Both the zone-reservation and progressive-ratio adaptation suggest that conditioned animals recognized the contingency between zone crossing and drug release, rather than responding solely on the basis of spatial preference or habit. Taken together, these findings further validate the operant nature of the paradigm and demonstrate that adult zebrafish can exert graded effort to obtain drug reinforcement.

We further demonstrated that preference for the drug-paired zone was driven by dopaminergic signalling and that conditioned fish exhibited behavioural signs consistent with withdrawal. Pharmacological disruption of dopamine signalling significantly reduced the choice ratio, suggesting that this engagement with the drug-paired zone relies on conserved reward circuitry.

In addition, the development of increased stress and anxiety following drug discontinuation is consistent with the development of withdrawal-like symptoms and suggests that repeated ethanol consumption has impacted the neurobiological functions of these animals, mirroring the adverse effects seen in both humans and animal models after drug discontinuation^55^. Human alcohol withdrawal is generally an acute syndrome lasting several days, with symptoms beginning 6–24 h after the last drink, peaking at 24–72 h, and resolving in most uncomplicated cases within 4–7 days^56^. Therefore, we believe that the observed effect in ethanol-conditioned zebrafish is in line with human withdrawal symptoms.

## Conclusion

In the future, it will be interesting to test whether this individual fish conditioning can be adapted to different drugs of abuse and to test other aspects of substance abuse disorder, such as extinction, relapse or incubation of craving.

This self-administration paradigm enables a refined investigation of addiction-like and drug-seeking–related behaviours in adult zebrafish by supporting multi-day training while eliminating social confounds inherent to group-based designs. The observed self-administration behaviour engages in conserved neurobiological pathways implicated in reinforcement and dependence across vertebrate species, including humans. As such, this assay provides a versatile platform for both genetic and pharmacological studies, with particular potential for small-molecule screening and functional validation of addiction-related genes, ultimately contributing to the identification of novel mechanisms and therapeutic targets for substance use disorders.

## Data statement

Data and custom tracking algorithms are available upon request.

## Authorship

L.M. Conceptualization, Methodology, Investigation, Formal analysis, Visualization, Writing – Original draft, Writing – Reviewing and editing.

L.G. Conceptualization, Methodology, Investigation

R.T.P. Conceptualization, Resources, Supervision, Funding acquisition, Project Supervision

G.D.B Conceptualization, Formal Analysis, Resources, Methodology, Software, Validation, Visualization, Data Curation, Supervision, Funding acquisition, Project Supervision, Writing – Original draft, Writing – Reviewing and editing.

## Acknowledgments

We would like to acknowledge the LARSEM and the animal care facility at CERVO for providing Zebrafish husbandry, laboratory space, and equipment to facilitate portions of this research.

## References

1. Research (Alcohol) | Canadian Centre on Substance Use and Addiction. https://www.ccsa.ca/en/guidance-tools-resources/substance-use-and-addiction/alcohol/research.

2. Sacks, J. J., Gonzales, K. R., Bouchery, E. E., Tomedi, L. E. & Brewer, R. D. 2010 National and State Costs of Excessive Alcohol Consumption. Am. J. Prev. Med. 49, e73–e79 (2015).

3. World Health Organization. Alcohol. https://www.who.int/news-room/fact-sheets/detail/alcohol (2024).

4. Understanding Alcohol Use Disorder | National Institute on Alcohol Abuse and Alcoholism (NIAAA). https://www.niaaa.nih.gov/publications/brochures-and-fact-sheets/understanding-alcohol-use-disorder?utm_source=chatgpt.com.

5. Koob, G. F. & Volkow, N. D. Neurocircuitry of Addiction. Neuropsychopharmacology 35, 217–238 (2009).

6. Koob, G. F. & Volkow, N. D. Neurobiology of addiction: a neurocircuitry analysis. Lancet Psychiatry 3, 760–773 (2016).

7. Nguyen, L.-C. et al. Predicting Relapse After Alcohol Use Disorder Treatment in a High-Risk Cohort: The Roles of Anhedonia and Smoking. J. Psychiatr. Res. 126, 1–7 (2020).

8. Yang, W., Singla, R., Maheshwari, O., Fontaine, C. J. & Gil-Mohapel, J. Alcohol Use Disorder: Neurobiology and Therapeutics. Biomedicines 10, 1192 (2022).

9. Canver, B. R., Newman, R. K. & Gomez, A. E. Alcohol Withdrawal Syndrome. in StatPearls (StatPearls Publishing, Treasure Island (FL), 2026).

10. Becker, H. C. Alcohol Dependence, Withdrawal, and Relapse. Alcohol Res. Health 31, 348–361 (2008).

11. Winslow, B. T., Onysko, M. & Hebert, M. Medications for Alcohol Use Disorder. Am. Fam. Physician 93, 457–465 (2016).

12. Bahji, A. et al. PHARMACOTHERAPIES FOR ADULTS WITH ALCOHOL USE DISORDERS: A SYSTEMATIC REVIEW AND NETWORK META-ANALYSIS. J. Addict. Med. 16, 630–638 (2022).

13. Kelly, J. F., Greene, M. C., Bergman, B. G., White, W. L. & Hoeppner, B. B. How Many Recovery Attempts Does it Take to Successfully Resolve an Alcohol or Drug Problem? Estimates and Correlates From a National Study of Recovering U.S. Adults. Alcohol. Clin. Exp. Res. 43, 1533–1544 (2019).

14. Mathur, P. & Guo, S. Use of zebrafish as a model to understand mechanisms of addiction and complex neurobehavioral phenotypes. Neurobiol. Dis. 40, 66–72 (2010).

15. Klee, E. W. et al. Zebrafish: a model for the study of addiction genetics. Hum. Genet. 131, 977–1008 (2011).

16. Schenk, S., Bossé, G. D. & Peterson, R. T. Zebrafish Models of Drug Addiction. In Psychobiological Issues in Substance Use and Misuse (Routledge, 2020).

17. Müller, T. E. et al. Understanding the neurobiological effects of drug abuse: Lessons from zebrafish models. Prog. Neuropsychopharmacol. Biol. Psychiatry 100, 109873 (2020).

18. Stewart, A. et al. Zebrafi sh models to study drug abuse-related phenotypes. 22, 95–105 (2011).

19. Lau, B., Bretaud, S., Huang, Y., Lin, E. & Guo, S. Dissociation of food and opiate preference by a genetic mutation in zebrafish. Genes Brain Behav. 5, 497–505 (2006).

20. Cousin, M. A. et al. Larval Zebrafish Model for FDA-Approved Drug Repositioning for Tobacco Dependence Treatment. PLoS ONE 9, e90467 (2014).

21. Kedikian, X., Faillace, M. P. & Bernabeu, R. Behavioral and Molecular Analysis of Nicotine-Conditioned Place Preference in Zebrafish. PLoS ONE 8, e69453 (2013).

22. Darland, T. & Dowling, J. E. Behavioral screening for cocaine sensitivity in mutagenized zebrafish. Proc. Natl. Acad. Sci. 98, 11691–11696 (2001).

23. Bretaud, S. et al. A choice behavior for morphine reveals experience-dependent drug preference and underlying neural substrates in developing larval zebrafish. Neuroscience 146, 1109–1116 (2007).

24. Petzold, A. M. et al. Nicotine response genetics in the zebrafish. Proc. Natl. Acad. Sci. 106, 18662–18667 (2009).

25. Bossé, G. D. & Peterson, R. T. Development of an opioid self-administration assay to study drug seeking in zebrafish. Behav. Brain Res. 335, 158–166 (2017).

26. Klee, E. W. et al. Zebrafish: a model for the study of addiction genetics. Hum. Genet. 131, 977–1008 (2012).

27. Kily, L. J. M. et al. Gene expression changes in a zebrafish model of drug dependency suggest conservation of neuro-adaptation pathways. J. Exp. Biol. 211, 1623–1634 (2008).

28. Tran, S. & Gerlai, R. Time-course of behavioural changes induced by ethanol in zebrafish (Danio rerio). Behav. Brain Res. 252, 204–213 (2013).

29. Tran, S., Chatterjee, D. & Gerlai, R. An integrative analysis of ethanol tolerance and withdrawal in zebrafish (Danio rerio). Behav. Brain Res. 276, 161–170 (2015).

30. Holcombe, A., Howorko, A., Powell, R. A., Schalomon, M. & Hamilton, T. J. Reversed Scototaxis during Withdrawal after Daily-Moderate, but Not Weekly-Binge, Administration of Ethanol in Zebrafish. PLOS ONE 8, e63319 (2013).

31. Martin-Fardon, R. & Weiss, F. Modeling Relapse in Animals. Curr. Top. Behav. Neurosci. 13, 403–432 (2013).

32. Vendruscolo, L. F. & Roberts, A. J. Operant alcohol self-administration in dependent rats: focus on the vapor model. Alcohol Fayettev. N 48, 277–286 (2014).

33. Holgate, J. Y., Shariff, M., Mu, E. W. H. & Bartlett, S. A Rat Drinking in the Dark Model for Studying Ethanol and Sucrose Consumption. Front. Behav. Neurosci. 11, (2017).

34. Huynh, N., Arabian, N. M., Asatryan, L. & Davies, D. L. Murine Drinking Models in the Development of Pharmacotherapies for Alcoholism: Drinking in the Dark and Two-bottle Choice. J. Vis. Exp. JoVE 10.3791/57027 (2019) doi:10.3791/57027.

35. Slosky, L. M. et al. Establishment of multi-stage intravenous self-administration paradigms in mice. Sci. Rep. 12, 21422 (2022).

36. Panlilio, L. V. & Goldberg, S. R. Self-administration of drugs in animals and humans as a model and an investigative tool. Addict. Abingdon Engl. 102, 1863–1870 (2007).

37. Kuhn, B. N., Kalivas, P. W. & Bobadilla, A.-C. Understanding addiction using animal models. Front. Behav. Neurosci. 13, 262 (2019).

38. Stefański, R. et al. Active versus passive cocaine administration: Differences in the neuroadaptive changes in the brain dopaminergic system. Brain Res. 1157, 1–10 (2007).

39. Sterling, M. E., Karatayev, O., Chang, G.-Q., Algava, D. B. & Leibowitz, S. F. Model of voluntary ethanol intake in zebrafish: Effect on behavior and hypothalamic orexigenic peptides. Behav. Brain Res. 278, 29–39 (2015).

40. Nathan, F. M. et al. Contingent stimulus delivery assay for zebrafish reveals a role for CCSER1 in alcohol preference. Addict. Biol. 27, e13126 (2022).

41. Raine, J., Kibat, C., Banerjee, T. D., Monteiro, A. & Mathuru, A. S. chrna3 Modulates Alcohol Response. J. Neurosci. 45, (2025).

42. Suriyampola, P. S. et al. Zebrafish Social Behavior in the Wild. Zebrafish 13, 1–8 (2016).

43. Caperaa, M. et al. Development of sensorimotor responses in larval zebrafish: A comparison between wild-type and GCaMP6s transgenic line. Behav. Brain Res. 481, 115412 (2025).

44. Bosse, G. D. et al. The 5α-reductase inhibitor finasteride reduces opioid self-administration in animal models of opioid use disorder. J. Clin. Invest. 131, (2021).

45. Czachowski, C. L. & Samson, H. H. Breakpoint determination and ethanol self-administration using an across-session progressive ratio procedure in the rat. Alcohol. Clin. Exp. Res. 23, 1580–1586 (1999).

46. Hodos, W. Progressive Ratio as a Measure of Reward Strength. Sci. N. Y. NY 134, 943–944 (1961).

47. Richardson, N. R. & Roberts, D. C. S. Progressive ratio schedules in drug self-administration studies in rats: a method to evaluate reinforcing efficacy. J. Neurosci. Methods 66, 1–11 (1996).

48. Sciascia, J. M., Mendoza, J. & Chaudhri, N. Blocking dopamine D1-like receptors attenuates context-induced renewal of Pavlovian-conditioned alcohol-seeking in rats. Alcohol. Clin. Exp. Res. 38, 418–427 (2014).

49. Tiglao, S. M., Meisenheimer, E. S. & Oh, R. C. Alcohol Withdrawal Syndrome: Outpatient Management. Am. Fam. Physician 104, 253–262 (2021).

50. Krook, J. T., Duperreault, E., Newton, D., Ross, M. S. & Hamilton, T. J. Repeated ethanol exposure increases anxiety-like behaviour in zebrafish during withdrawal. PeerJ 7, e6551 (2019).

51. Mathur, P. & Guo, S. Differences of Acute versus Chronic Ethanol Exposure on Anxiety-Like Behavioral Responses in Zebrafish. Behav. Brain Res. 219, 234–239 (2011).

52. Egan, R. J. et al. Understanding behavioral and physiological phenotypes of stress and anxiety in zebrafish. Behav. Brain Res. 205, 38–44 (2009).

53. Sanchez-Roige, S., Kember, R. L. & Agrawal, A. Substance use and common contributors to morbidity: A genetics perspective. eBioMedicine 83, 104212 (2022).

54. Deak, J. D. & Johnson, E. C. Genetics of substance use disorders: a review. Psychol. Med. 51, 2189–2200.

55. Jesse, S. et al. Alcohol withdrawal syndrome: mechanisms, manifestations, and management. Acta Neurol. Scand. 135, 4–16 (2017).

56. Jesse, S. et al. Alcohol withdrawal syndrome: mechanisms, manifestations, and management. Acta Neurol. Scand. 135, 4–16 (2017).

